# Meta-control of the exploration-exploitation dilemma emerges from probabilistic inference over a hierarchy of time scales

**DOI:** 10.1101/847566

**Authors:** Dimitrije Marković, Thomas Goschke, Stefan J. Kiebel

**Affiliations:** Chair of Neuroimaging, Faculty of Psychology, Technische Universität Dresden, 01062 Dresden, Germany; Chair of General Psychology, Faculty of Psychology, Technische Universität Dresden, 01062 Dresden, Germany; The Centre for Tactile Internet with Human-in-the-Loop (CeTI), Technische Universität Dresden, 01062 Dresden, Germany

**Keywords:** meta-control, arbitration, exploration-exploitation dilemma, hierarchy of time scales, probabilistic inference, prior over policies

## Abstract

Cognitive control is typically understood as a set of mechanisms which enable humans to reach goals that require integrating the consequences of actions over longer time scales. Importantly, using routine beheavior or making choices beneficial only at a short time scales would prevent one from attaining these goals. During the past two decades, researchers have proposed various computational cognitive models that successfully account for behaviour related to cognitive control in a wide range of laboratory tasks. As humans operate in a dynamic and uncertain environment, making elaborate plans and integrating experience over multiple time scales is computationally expensive, the specific question of how uncertain consequences at different time scales are integrated into adaptive decisions remains poorly understood. Here, we propose that precisely the problem of integrating experience and forming elaborate plans over multiple time scales is a key component for better understanding how human agents solve cognitive control dilemmas such as the exploration-exploitation dilemma. In support of this conjecture, we present a computational model of probabilistic inference over hidden states and actions, which are represented as a hierarchy of time scales. Simulations of goal-reaching agents instantiating the model in an uncertain and dynamic task environment show how the exploration-exploitation dilemma may be solved by inferring meta-control states which adapt behaviour to changing contexts.

## Introduction

The concept of cognitive control is generally used as a summary term for a set of processes that enable humans to flexibly configure perceptual, emotional, and response selection processes in accordance with superordinate goals. These processes are especially pronounced when goal attainment requires novel or non-routine action sequences, and there is competition from otherwise stronger habitual or impulsive responses (Botvinick and Cohen 2014; Egner 2017; Goschke 2003, 2013; Miller and Cohen 2001). Cognitive control is considered essential for some of the most advanced cognitive capacities of humans, such as the ability to pursue long-term goals and to respond flexibly to changing contexts and task demands. However, much of the experimental research on cognitive control has focused on relatively simple laboratory tasks, as, for instance, interference paradigms such as Stroop or flanker task (e.g., Kalanthroff et al. 2018; Scherbaum et al. 2011), or paradigms assessing cognitive flexibility such as task switching (Koch et al. 2018). Many of these tasks are aimed at inducing conflicting internal representations, which trigger responses that are in contradiction to the instructed task goal and may lead to an incorrect response. Such tasks have been remarkably useful as psychological ‘probes’ into component mechanisms of cognitive control such as response inhibition or goal shielding, as they enable researchers to study how the brain copes with crosstalk between conflicting representations and competing responses. Accordingly, many computational models of cognitive control postulate a hierarchical mechanism, where higher-level representations of goals or task-sets serve as a biasing signal, which modulates processing at a lower level, such that information congruent with instructed goals gains higher priority in determining the selection of responses (Cohen 2017; Goschke 2003, 2013; Miller and Cohen 2001; Scherbaum et al. 2012). More recently, hierarchical models have been used to establish how the brain might determine the intensity and allocation of biasing signals to specific tasks, based on the estimated costs and benefits of recruitment of control (e.g., Shenhav, Botvinick, and Cohen 2013).

Although these approaches to study and model cognitive control have been highly successful and are widely used, they focus on a specific class of task, which differ in a key aspect from real-life goal-reaching scenarios in which humans typically use cognitive control. This difference is that experimental cognitive control tasks typically require little planning, i.e. the participant is not required to plan ahead (e.g. across several trials) to choose an action. By planning we mean that to select an action, an agent has to predict the consequences of this action over longer time periods than just the current trial in a task. Clearly, planning is an important part of cognitive control because it is necessary to reach goals, as formalized mathematically in the reinforcement learning and active inference frameworks (Botvinick, Niv, and Barto 2009; Friston et al. 2017; Pezzulo, Rigoli, and Friston 2015). Some recent studies have tapped explicitly into planning using sequential decision making tasks, where, to reach a goal over a series of trials, participants have to plan ahead for around 30 seconds, (e.g., Economides et al. 2014; Kolling, Wittmann, and Rushworth 2014; Schwartenbeck et al. 2015).

### Planning in uncertain environments

Although not always obvious to us, human planning is for many tasks in daily life a computational feat yet unrivalled by any machine. Research in robotics and artificial intelligence has found that planning ahead, in an online fashion, in our typically uncertain environment is a hard problem for artificial agents, (e.g., Kurniawati et al. 2011). Even for mundane activities such as safely driving a car through typical traffic, artificial planning performance is currently well below human routine performance (for a current review see Schwarting, Alonso-Mora, and Rus 2018). Here, planning is required because a car responds rather slowly to one’s actions so that one must predict the consequences of one’s own actions into the future for at least a few seconds or even longer, especially in the presence of other traffic participants, whose behaviour must also be predicted. Although there are recent findings that artificial agents perform better than humans in specific planning tasks like playing the board game Go (Silver et al. 2017), the question is what makes planning challenging in scenarios such as driving a car. Here, we will focus on two of these features, which are also probably the most relevant for addressing cognitive control research questions.

Firstly, for a goal-directed agent, most environments are packed with uncertainty. This uncertainty is induced by various sources. For example, the noise of exteroceptive and interoceptive input, the usually hidden causes of events or the intentions of other agents that must be inferred from their behaviour. In a board game like Go, the only source of uncertainty is the opponent, whereas the agent has full knowledge of the current state of the board, which is free from sensory noise and contains no hidden parts, and the rules are deterministic and define unambiguously which moves are admissible and which are forbidden. This stands in stark contrast to real-life planning scenarios like driving, where we cannot observe all other traffic participants continuously with high precision, objects may be blocked from our view, traffic rules are probabilistic, because other participants may violate them, and instead of a single opponent, there are multiple traffic participants with hidden intentions. These sources of uncertainty in real environments make planning difficult because the number of possible ways in which the environment may develop grows massively the further into the future one tries to plan ahead (Huys et al. 2012).

Secondly, in our environment, things change at different time scales. For example, in the board game Go the relevant time scale is well defined and choices matter only within the confines of the game, similar to a trial or a block of trials in an experiment. In our environment, very different time scales co-exist. For instance, a pedestrian’s quick glance over her shoulder (which occurs on a time scale of a few hundred milliseconds) may indicate that she will be crossing the street (which may take several seconds), which may be part of the action plan to meet a friend at a café (which may span a time scale of two hours), which may be motivated by the intention to maintain positive social relations (which spans a time scale of years), (e.g., Mylopoulos and Pacherie 2019). In other words, in real life situations we are confronted with uncertainty about the relevance of different time scales, that is, one problem is to infer the relevant time scales for one’s planning and goal reaching. Although it is clear that Go strategies evolve also over several time scales (a single move, several moves, the whole game), shorter time scales or time scales beyond the end of the game are typically not relevant for an agent playing Go. In contrast, in real-life an observed quick glance can provide rich information for slower, more coarse-grained time scales. Similarly, very slow time scales are highly relevant as real life hopefully lasts for many years to come.

There is recent experimental and theoretical evidence in the cognitive neurosciences that these multiple time scales are a critical dimension of how the brain structures its environment (Badre and Nee 2018; Chaudhuri et al. 2015; Dixon and Christoff 2017; Kiebel, Daunizeau, and Friston 2008; Koechlin, Ody, and Kouneiher 2003). In the domain of cognitive control, the relevance of different time scales is well established in the context of, for instance, intertemporal choice conflicts, where agents have to choose between a smaller reward that can be obtained immediately versus a larger reward that can only be obtained only after a delay (Dai, Pleskac, and Pachur 2018; Kable 2014; Scherbaum et al. 2013).

### Uncertainty and a hierarchy of time scales

Below we will present a simple experimental task that requires planning at two different time scales under several sources of uncertainty. One of the aims of this paper is to illustrate how one can build a computational agent for this task and implement the function of cognitive control in this mechanistic model. The conceptual backbone of the model is that the representation of environmental dynamics is organized as a hierarchy of time scales (Kiebel, Daunizeau, and Friston 2008). Such modelling approaches have been proposed in cognitive control in the context of hierarchical reinforcement learning (HRL), (e.g., Botvinick and Weinstein 2014; Holroyd and McClure 2015) and are naturally also an increasingly relevant topic in artificial intelligence research (e.g., Bacon and Precup 2018; Pang et al. 2019; Le, Vien, and Chung 2018; Mnih et al. 2015). In general, HRL models are based on the idea that action sequences can be chunked and represented as a new temporally extended state, (see also Maisto, Donnarumma, and Pezzulo 2015) for a probabilistic modelling alternative. For example, making tea is a state that lasts about 30 seconds and requires performing a series of actions. Each of these actions (e.g. to get some water) is at a faster, more fine-grained time scale and last only a few seconds. This principled idea to represent behaviour as a hierarchy of sequences has also been proposed as a way how one may understand recent findings in fields such as speech (Hasson et al. 2008), memory and the hippocampus (Collin, Milivojevic, and Doeller 2017), and decision making (Hunt and Hayden 2017). Note that the principled idea that goal-directed control is organised as a hierarchy with elements represented at different time scales can be traced back to concepts outlined for example by Miller, Galanter, and Pribram (1960) and pursued in action control theories (Gollwitzer and Bargh 1996; Heckhausen and Kuhl 1985; Kuhl and Goschke 1994). We will use the principle as exemplified by recent HRL modelling work but critically complement the resulting model by three components, which we believe are important to explain specific cognitive control phenomena. Note that all three components have been used before in probabilistic modelling approaches and are not novel by themselves. Our point is that the combination of these specific model features may make a difference for research into cognitive control.

Firstly, as motivated above, planning in our environment must incorporate various sources of uncertainty, which requires that we formulate the hierarchical model probabilistically (see Methods for details). Secondly, hierarchical reinforcement learning models previously applied in the cognitive neurosciences (e.g., Holroyd and McClure 2015) typically assume that agents aim at maximizing future return (instrumental value - IV). This approach works well for modelling and analysing experimental tasks, which require participants to reach goals in an already well-learned task environment. However, when considering cases in which an agent has not yet learned its task environment, actions should not only serve the maximization of reward but also the reduction of uncertainty about task-relevant states and parameters (Ghavamzadeh 2015). To be able to model such uncertainty-reducing, explorative actions of an agent, we will use the expected free energy, which combines instrumental value with the epistemic value of different actions, thereby leading to a reduction of uncertainty about the state of the world (Kaplan and Friston 2018). Thirdly and most importantly, we introduce specific hidden states, which we call in the following ‘meta-control states’. We use these states for letting the agent represent which policies it should prefer for planning. Meta-control states do not represent the environment but represent how the agent should behave in a specific context. As we will show below meta-control states can be inferred by the agent online and be used to provide a learnable mapping from the task context to the subset of behavioural policies that are most suitable for reaching a goal. We will also show that these meta-control states effectively cause the computation of control signals, which guide concrete low-level behaviour. In simulations, we will focus on the usefulness of meta-control states to solve the exploration-exploitation dilemma and will discuss how inference over meta-control states may be used to resolve other cognitive control dilemmas.

### Cognitive control dilemmas

Agents with an extended future time perspective, who pursue goal-directed action in changing and uncertain environments are confronted with a set of antagonistic adaptive challenges. These challenges can be conceived of as fundamental *control dilemmas*, which require a context-sensitive adjustment of complementary control modes and control parameters (Goschke 2003, 2013; Goschke and Bolte 2014). For instance, while the ability to shield long-term goals from competing responses promotes behavioural stability and persistence, it increases the risk of overlooking potentially significant changes in the environment and may lead to rigid and perseverative behaviour. Conversely, while a broad scope of attention supports background-monitoring for potentially significant changes and facilitates flexible goal switching, it also increases distractibility and may lead to volatile behaviour that is driven by every minor change in the environment (Dreisbach and Goschke 2004; Goschke and Bolte 2014). Agents must thus not only decide which action is best suited to attain a goal, but they have to cope with meta-control problems (e.g., should one ignore an unexpected change and shield a current goal from distraction or should one process task-irrelevant information, because it may signal that one should switch to a different goal?). Given that antagonistic adaptive constraints cannot be satisfied simultaneously to an arbitrary degree, because stable versus flexible control modes incur complementary costs and benefits, goal-directed agents must solve *meta-control problems*, which raise the question how the brain achieves a context-sensitive balance between complementary control modes and how control parameters are adjusted to optimize goal attainment in changing and uncertain environments.

While control dilemmas arise in a range of processing domains (e.g., goal shielding vs. goal shifting; focused attention vs. background-monitoring; anticipation of future needs vs. responding to current desires; computationally demanding but flexible goal-directed control vs. less demanding but inflexible habitual control, see below for a brief discussion), here we focus on the trade-off between exploration and exploitation as one of the most widely investigated control dilemmas (Blanchard and Gershman 2018; Cohen, McClure, and Yu 2007; Addicott et al. 2017).

It is obviously adaptive for agents to exploit and select those actions that maximized reward in the past. However, to learn about such actions or find better ones, agents must explore previously untried actions. Thus exploitation may prevent learning about task-relevant actions and states; conversely, exploration supports learning and may return relatively little reward or even lead to risky behaviour.

In the following, we will describe a simple task incorporating planning under uncertainty in the presence of two time scales and an agent that can perform adaptively in this task. Our focus will be on using meta-control states to describe how agents can adapt their behaviour in a way that is reminiscent of the exertion of cognitive control.

### Simple experimental task

We use a sequential decision making task, similar to previous studies where participants had to collect points in a series of trials to surpass a known point threshold (e.g., Kolling, Wittmann, and Rushworth 2014). The task combines the rationale of such sequential decision making tasks with aspects of probabilistic reversal learning tasks (Cuevas Rivera et al. 2018; Markovic, Reiter, and Kiebel 2019). Instead of using only two different contexts, our task comprises six contexts. The goal of the design is to differentiate unambiguously between explorative and exploitative behaviour of the agent.

In the task, runs of five trials form a segment, during which the participant can collect points in each trial by choosing one of four different options. Each of these options returns probabilistically one blue point, one red point, or no point. The number of collected points is evaluated after the fifth trial, where the reward is only given if the agent succeeded to collect at least four points of the same colour. For example, 4 red points and 0 blue points are rewarded, while 3 red points and 1 blue point are not rewarded. Although this setup and the following task description may appear quite complex in relation to typical cognitive control tasks like the Stroop task, we found that this level of task complexity is required to measure clear behavioural differences when doing the task in either an explorative or exploitative mode.

The experiment consists of a series of five-trial segments where, in addition, we impose changes at a slower time scale to introduce so-called contexts. A context determines the payoff matrix for all five trials of a segment, i.e. a context determines the point outcomes and probabilities for the four options. Context changes occur whenever five segments, i.e. 25 trials, have been completed. Similar to a typical reversal learning task, changes are not explicitly indicated so that the agent can infer the current context only from a sequence of choice outcomes. There are six different contexts (see Figure 1), where the conceptual idea of the experiment is that in three of these contexts explorative behaviour is more successful than exploitative behaviour, and vice versa in the other three contexts. This means that a goal-directed agent, which employs meta-control, should use either explorative or exploitative behaviour depending on the context. The six contexts come in three pairs. Each context pair, e.g. context 1A and 1B (see Figure 1), consists of the context variant A in which exploitative behaviour should be preferred, and the very similar context variant B in which explorative behaviour should be preferred. As can be seen in Figure 1, for each context pair, the variants A and B differ only in the payoff of one of the four choice options while the payoffs of the remaining three options are identical. For example, for both contexts 3A and 3B, option 1 returns a red point with 80% probability, and options 2 and 3 return a red point with 10% probability each. The one different option is number 4, where in variant B a red point is received with 100% probability but in variant A with 0% probability. This specific construction of context pairs has the effect that if an agent knows that the current context is context 3 but does not know its variant (A or B), option 1 has the highest expected reward (0.8 red points) of all options while the expected reward for option 4 is only 0.5 red points. This makes explorative versus exploitative behaviour easily identifiable because an exploitative agent, once it infers the context pair, e.g. number 3, will try to maximize expected reward by choosing the 80% option number 1, while an explorative agent would reduce its uncertainty about the context variant (A or B) by choosing option 4. As in real life, sometimes exploration pays off, and if an agent with explorative behaviour finds itself in one of the three context variants B, it will outperform an agent with exploitative behaviour because the explorative agent will quickly find the 100% option. However, in the three context variants A, an agent with exploitative behaviour will collect on average more reward than an explorative agent because it sticks with the 80% option.

**Figure 1:**
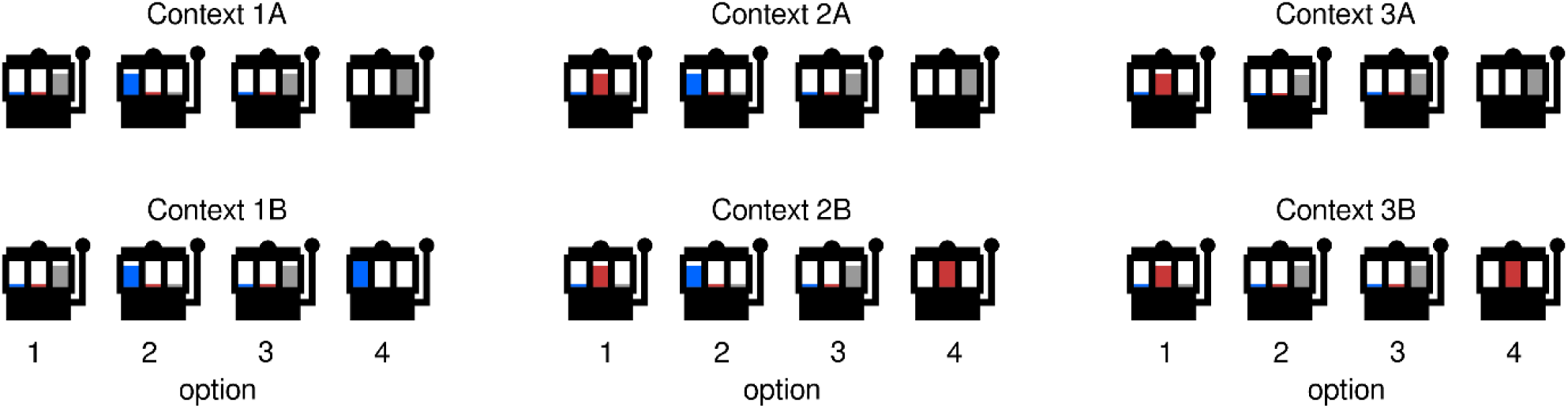
Illustration of payoff in six different contexts. Each context is defined by the payoff probabilities associated with four different options. In context variants A (top row), agents with exploitative behaviour will on average be more successful in reaching the segment-wise goal to surpass the threshold of four points of a single colour, in comparison to agents with explorative behaviour, and vice versa in the context variants B (bottom row). Furthermore, the only difference between each context pair, e.g. contexts 1A and 1B, is the option with the point probability of 100% in the B variants, i.e. option 4 in each context. All other options return a blue, red or no point with 80% probability and the other two outcomes with 10% probability. Note that option types (point probabilities associated with an option) are shared across context, e.g. the point probabilities (80% blue point, 10% red point, 10% no point) is used four times in contexts 1A, 1B, 2A and 2B. If an agent does not know the current context variant (A or B), the expected return of choosing the fourth option is lower compared to options associated with 80% point probability, e.g. option 1 in context pair 3A/3B. However, options which return a point (or no point) with 100% probability are the most informative because they resolve the uncertainty about the context variant A or B.

The main point of the simulations below will be to demonstrate that, given the task, we can now build a probabilistic inference agent that changes its exploration-exploitation behaviour depending on context. We will do this by using a model based on a hierarchy of time scales and active inference, where the agent does not only performe inference over hidden context and meta-control states but also inference over control signals which determine preferable modes of behaviour. As we will show below in detail, agents doing the task will learn task parameters during a training period, just as human participants would do. Specifically, agents have to learn the outcome probabilities (blue, red, no point) associated with each option, in each of the six contexts to be successful in the task. Importantly, the agent is informed that there are only six different contexts.

### Behavioural model

We constructed the task such that the current context can be inferred only with uncertainty due to the probabilistic outcomes of the four options. Consequently, we will model decision making by the agent as a partially observable Markov decision process (POMDP), (Littman 2009; Kaelbling, Littman, and Cassandra 1998). As the agent cannot directly observe the underlying states, e.g. which of the six contexts is the current one, the agent has to form beliefs over possible states and make decisions based on these beliefs. This means that the decisions of the agent are made under uncertainty about the current context. To build an agent and reflect the task structure of trials embedded into segments under specific contexts, we first define a generative hierarchical model with two levels. This generative model defines a set of rules and statistical dependencies that an agent uses to make probabilistic predictions and infer its belief about the underlying state of the environment from choice outcomes.

Specifically, the agent’s generative model represents the probabilistic mapping between the four choices and the possible outcomes of receiving a point. The agent represents the duration of each segment (five trials) and that success depends on collecting at least four coloured points of a single colour. The agent represents six possible contexts at the second level, i.e. it is informed that in each trial one of the six contexts is active. The agent is not informed that the context switches every five segments but has the knowledge that the context can change to any other context with probability *p=1/5* between segments. Note that in all simulations below the agent has no expectation about the identity of the next context and no means of learning such expectations. See Figure 2 for a graphical representation of the two-level hierarchical model. The importance of the second (higher) level is that the agent represents at this level the slow time scale of segments and possible context switches between segments. Each of the six contexts is associated with a specific probabilistic choice-outcome mapping, see Figure 1. This mapping is used at the lower level, which represents the trial-specific faster time scale of choosing options and observing outcomes. The agent will be initially uninformed about these context-specific choice-outcome mappings and has to learn these mappings in a training period by interacting with the task environment.

**Figure 2.**
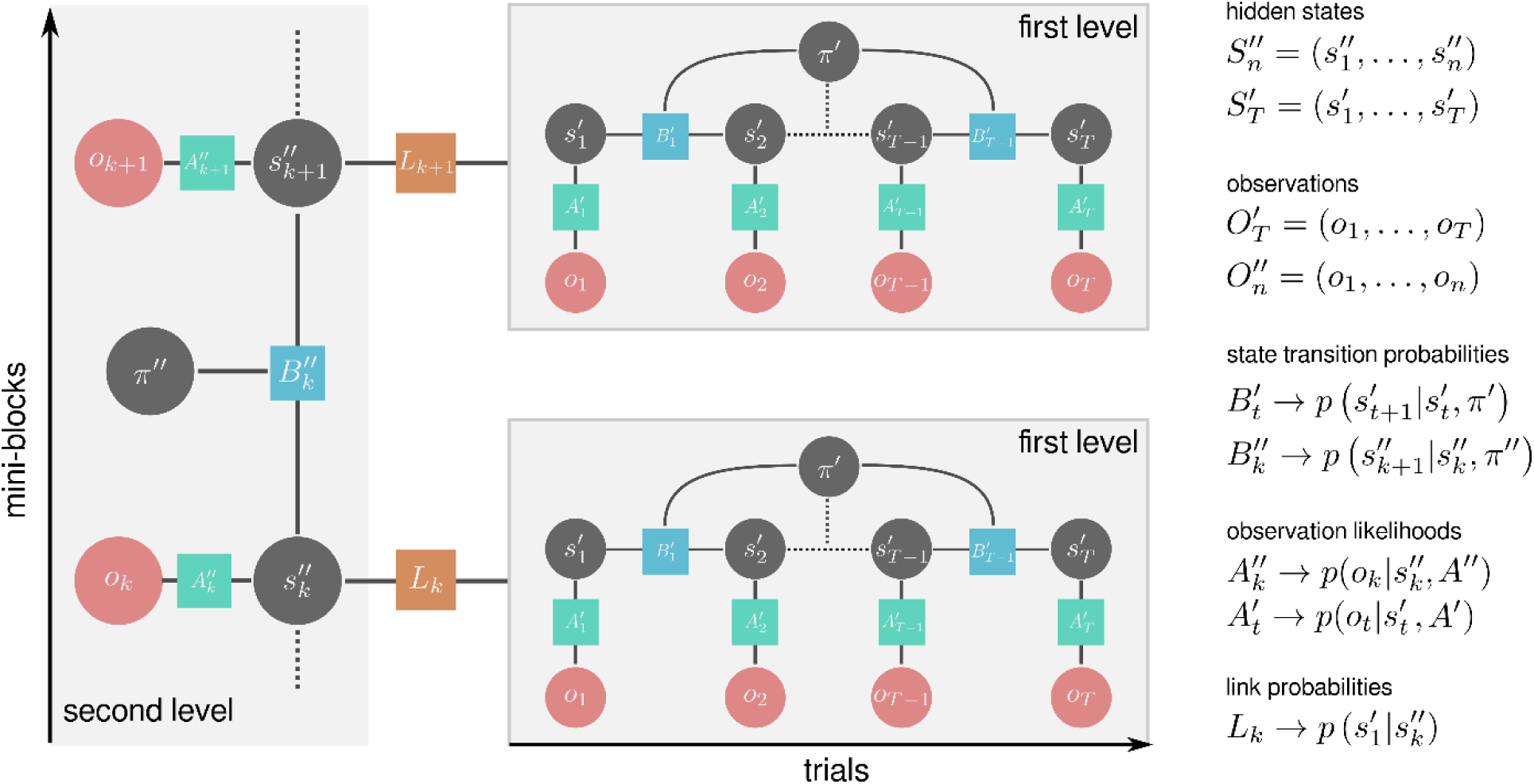
Factor graph representation of the hierarchical generative model for the presented task. The graph consists of two types of nodes: (i) Random variables (circles), which can be either evidence variables (red) whose value is observed or hidden state variables (grey) whose value has to be inferred. (ii) Factors (squares), which define the relationship between random variables. At the highest level of the hierarchy, the agent entertains beliefs (a probability distribution over the set of possible states) about the current context and its meta-control state, hence 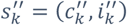, the c-i pair defines the observation likelihood of the outcome o_k_ at the end of a segment (success or failure). The behavioural policy at the second level of the hierarchy π″ consists of selecting the appropriate meta-control state for the next segment, depending on the expected changes in the context. The link probability L_k_ relates second level states to the prior beliefs about the lower level states 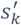. The lower level states factorise into the chosen options 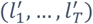 and auxiliary context and control states 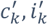 (fixed states during each segment) which capture lower level information about higher level states. Importantly, the auxiliary context states 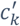 determine currently active observation likelihood, and the auxiliary control states 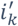 set prior over policies 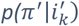 at the first level of the hierarchy. For details see the methods section.

To reach the segment-wise goal of collecting four points of a specific colour, the agent has to plan ahead and select behavioural policies, i.e. sequences of actions. When the agent has learned the relation between states (e.g., having two red and one blue point on trial 3 and being in context 2), and how a specific choice produces an outcome that will change this state (e.g., choosing option 2 may provide an additional blue point), the agent can make predictions about the consequences of selecting a specific policy, reaching further than a single trial.

To enable the agent to deal with the exploration-exploitation dilemma based on these predicted consequences, we will provide the agent with two aims: (i) to maximize the expected instrumental value, i.e., the amount of reward, here the number of segments, in which the agent has collected four points of one colour after the fifth trial of a segment and (ii) to maximize so-called epistemic value, the amount of information (see e.g., Kaplan and Friston 2018). As outlined in the previous section, we have designed our task such that the 4^th^ option in each context (see Figure 1) carries the highest epistemic value because it clearly differentiates between the context variants A or B, while its expected instrumental value is relatively low, in comparison to the 80% options. Note that pursuing both these aims is equivalent to minimising the expected free energy of an agent’s actions. In the task, the expected free energy is minimized by making choices which (i) brings the agent into a preferred state (obtaining a reward at the end of a segment) and/or (ii) reduces the agent’s uncertainty about hidden states (e.g., current context and choice-outcome probability).

## Simulations

Here we expose the agent to the task, see Figure 1 and the previous section for a description. We will proceed in three stages. First, to illustrate the basic features of the model, we will show the behaviour of agents that are fixed in their explorative versus exploitative stance, i.e. do not have meta-control. Second, we will introduce meta-control states, which enables an agent to resolve the exploration-exploitation dilemma by adapting its meta-control states in a context-dependent fashion. Third, we will show that the proposed model also enables an agent to infer that it should change its meta-control state already before a context switch when the agent can predict probabilistically an impending context change.

In the first illustrative simulation, we exposed agents to the task for 200 segments, i.e. 1,000 trials. In Figure 3, we show group mean success rates of three different agent types, where each group consists of *n* = 100 agents of the same type. One of these agents simply serves as a reference random choice agent. The other two agent types differ in their policy selection objective. In one case, the policy selection objective corresponds to the instrumental value (IV) only and in the other case to the expected free energy (EFE), i.e. the combined instrumental and epistemic value (see Priors over policies – expected free energy sub-section in Methods for details). In the task, maximizing IV only results in exploitative behaviour of an IV agent while an EFE agent is expected to show more explorative behaviour because of the EFE’s epistemic value component. We assume that the two agents have sufficiently learned the choice-outcome probabilities for the six contexts after 100 segments. Note that we used the alternating pattern of context variants A and B to maximize the need for adapting to a new context, see also below. As expected, there are large performance differences between context types A (indicated by black circles) and B. This is because in context types B, for each of the three contexts, there is the 4^th^ option that returns a point with 100% probability, see the task description above. In context variants B, the EFE agent reaches in context B types the highest performances because the affinity toward informative choices enables the agent not only to resolve the uncertainty about the current context but also to collect points with maximal probability. In context A types, the EFE agent has clearly a worse performance than the IV agent.

**Figure 3.**
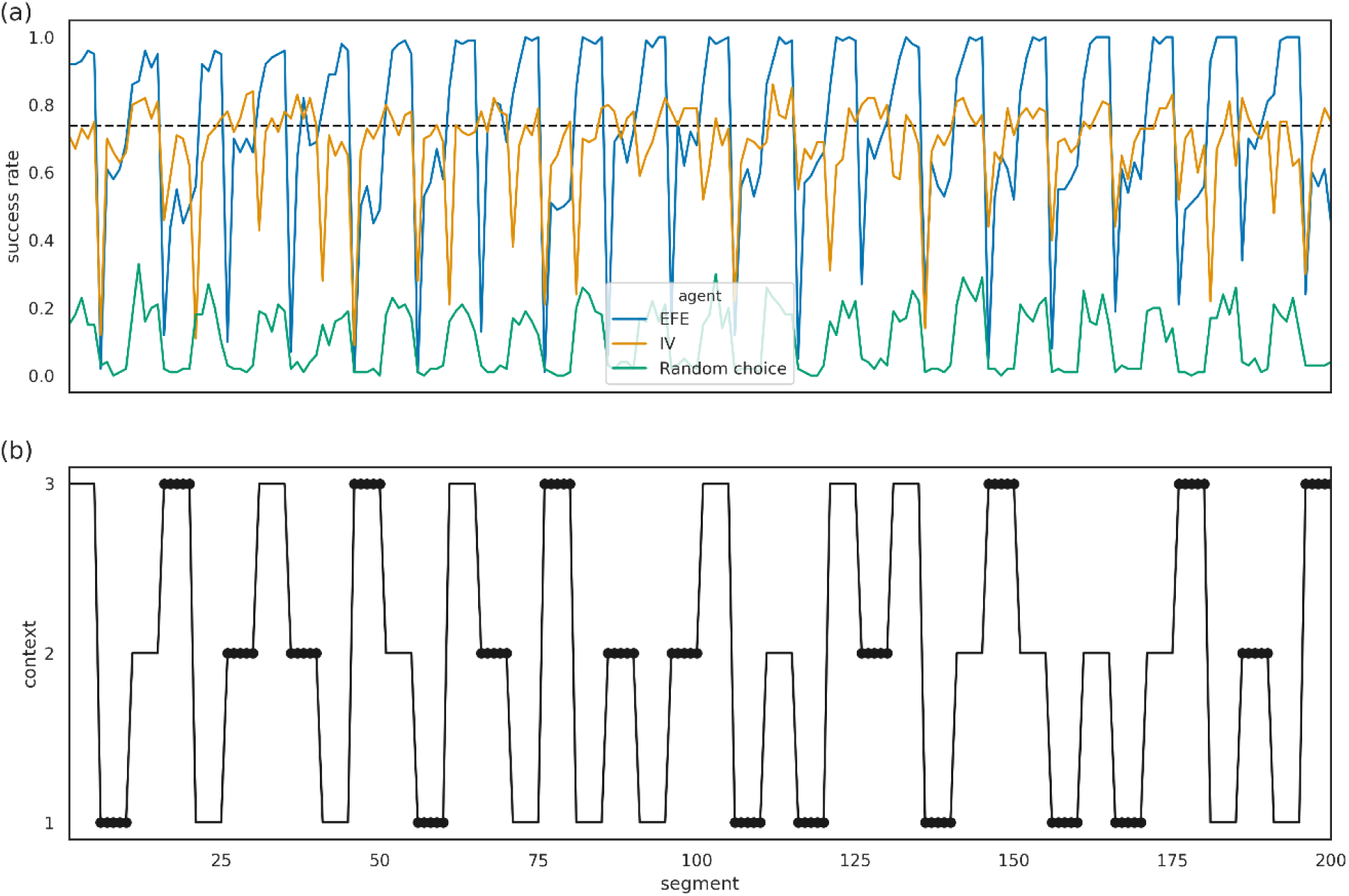
Success rates of three different agents. **(a)** Group mean success rates for the expected free-energy agent (EFE; blue line), instrumental agent (IV; orange line), and a random choice agent (green line), which randomly selects one of the four options on a trial with equal probabilities. The black dashed line denotes the expected success rate for always selecting an option which returns a coloured point with probability p = 0.8. **(b)** Context change schedule across segments. Circles denote segments under context variants A, in which exploration lowers success probability.

To understand the difference of mean success rates of the IV and EFE agents in both contexts variants A and B we now take a closer look at their choice probabilities. In Figure 4, one can see that the EFE agent is more likely to select the 4^th^ option, which is the most informative about the current context, see Figure 1. This allows the agent to resolve uncertainty about the context rapidly, leading to higher performance in context variants B and similar performance in context variants A as only a few trials are needed to resolve uncertainty and identify the true context.

**Figure 4.**
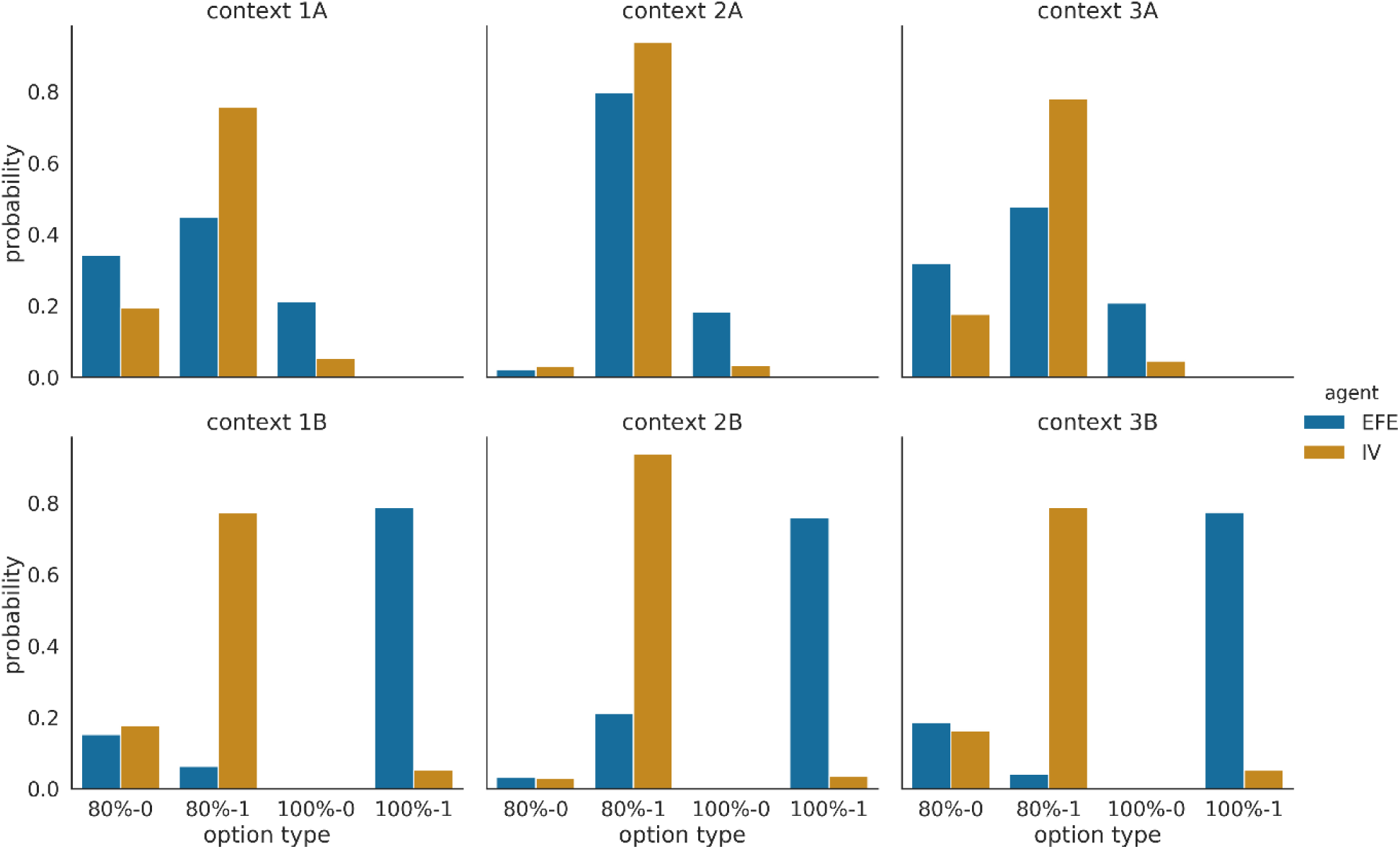
Probability of selecting different options in different contexts and context variants. The probabilities are estimated from 100 simulations for each agent type (the same as used for Figure 3) and pooled across the last 100 segments of the experiment. The three context variants A are shown in the upper row and the three context variants B in the lower row. The EFE agent (blue bars) selects the informative options (option type 100%-1) with the highest rate when exposed to variant B, and is also more likely to select the informative option (option type 100%-0) when exposed to variant A, compared to the IV agent (orange bars). For visualisation, we have pooled options that return a point with high probability (independent of the colour, 80%-1 and 100% −1) and options that return no points with high probability (80%-0 and 100%-0)

### Adaptive control of exploration-exploitation dilemma

Up to now, we have shown that there are interesting behavioural differences between an agent that just maximizes instrumental value (IV) and an expected free-energy (EFE) agent that, in addition, also considers information gain when selecting its policy. As we have found, not unexpectedly, the EFE agent follows more informative policies, which results, due to the task design, in a performance advantage in context variants B, and loss of performance in context variants A. Critically, the relative contributions of the instrumental and epistemic value to the policy selection were fixed in both the IV and EFE agent. However, one could argue that agents should be able to adapt their behavioural mode depending on the context, i.e. use autonomously controlled contributions of the two value terms for policy selection, akin to human meta-control.

Here, we implemented the conceptual idea to enable such meta-control in an agent by linking the inference over meta-control states, which define contributions of the instrumental and epistemic values, to policy selection. These meta-control states 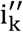 are part of the second level states 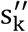 (see the graphical model in Figure 2) and linked to each context via observations of success or failure in each segment. Specifically, the meta-control states adapt the selection of policies by changing the prior over policies at the lower level (where we define this prior as the expected free energy), see also (Parr and Friston 2019). Intuitively, the prior over policies can be interpreted as a behavioural mode or a strategy because the prior simply tells an agent which action sequences it should currently prefer. Importantly, the prior over policies, depending on the meta-contol state, will either take the epistemic value term into account or ignore it. However, the uncertainty over currently preferred meta-control states will lead to a continuous weighting of the epistemic value term. The adaptive weighting biases the set of a priori viable policies which in turn influences the computations of the posterior over policies. We anticipate that such an adaptive agent will learn to be biased towards exploitative behaviour in context variant A and towards explorative behaviour in context variant B. In other words, an observer of the agent’s behaviour would possibly conclude that this agent resolves the exploration-exploitation dilemma by exerting meta-control.

Critically, the meta-control states do not represent external states of the environment but rather internal modes of behaviour. Note that the prior over policies does not exclude any policies in a hard-wired fashion. Rather, some policies become more likely to be selected than others.

To show this, we will compare the behaviour of this adaptive agent to the behaviour of the IV and EFE agents, which we used in the simulations above. These two non-adaptive agents represent the two extreme modes of the adaptive agent: the IV agent corresponds to a zero weighting of the epistemic value term, and the EFE agent to the unit weighting of the epistemic value term. In Figure 5a, we show the group mean success rates of the adaptive and the two non-adaptive agents, using the same task design, as shown in Figure 3b. One can see that the adaptive agent is on average similar in performance to the explorative agent in the context variants B, which shows that the adaptive agent switches to an exploratory mode if in contexts where maximum success rate can be obtained using exploratory behaviour. However, in context variants A, the performance of the adaptive agent is only slightly better as compared to the exploratory agent and far below the exploitative agent. The reason for this apparent non-adaptation to an exploitatory mode can be seen in Figure 5b, where we plotted the trajectories of the weighting 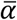 of the epistemic value for policy evaluation, i.e. a value of 1 indicates that the adaptive agent is in an explorative mode, while a value of 0 indicates an exploitative mode. Due to the learning in the first half of the experiment, the dynamics of the weighting factor 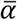 are history dependent, as can be seen for the trajectories of 100 agent instances doing exactly the same task with the same context sequence but with differently sampled outcomes, see Figure 5b (blue lines). This implies that the stochasticity of the outcomes interacts with the learning process on both levels of the hierarchy generating unique, adaptive behaviour that is sensitive to previous experience. To further quantify the differences between the adaptive agent and the two non-adaptive agents, we looked at two other quantities: (i) The context inference accuracy, see **Figure 6**a. We have defined context inference accuracy as the probability that the agent correctly identifies the current context (measured by the highest posterior probability for the true context). The adaptive agent achieves high levels of inference accuracy in both context variants. In other words, the adaptation of the behavioural modes does not have a detrimental impact on the ability of the adaptive agent to resolve its uncertainty about the current state of the world. (ii) The success probability of different agents and their time course as shown in **Figure 6**b (see Methods for the precise definition of the success probability). Note that unlike the success rate in Figure 5a, which is computed as a mean over multiple agent instances, the success probability is agent-instance specific, i.e. specific to a single agent. In context variants B, the success probability of the adaptive agent is as high as the success probability of the exploratory agent. However, in context variants A, the adaptive agent’s success probability is lower as compared to the one of the exploitative agent, but significantly higher than the explorative agent (p<0.05 as per Wilcoxon signed-rank test for all relative segment values). This average, lower performance can be directly related to the wide distribution of trajectories of the weighting factor 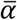 as shown in Figure 5b. In other words, many out of the 100 adaptive agents, due to the high stochasticity of the task (e.g., there is uncertainty on the current context and on the actual reward probabilities), do not learn how to behave exploitatively in context variants A. This point of variability in experience-dependent adaptation is stressed by showing the average success probability of a subset of ten instances of the adaptive agent which learned to down-regulate 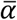. We selected these ten agent instances using the criterion of a downregulated epistemic weight below the 0.5 level in context variants A. One can clearly see (**Figure 6**b, black line) that the average success probability of this subset of adaptive agents is close to the performance level of the exploitative agent.

**Figure 5.**
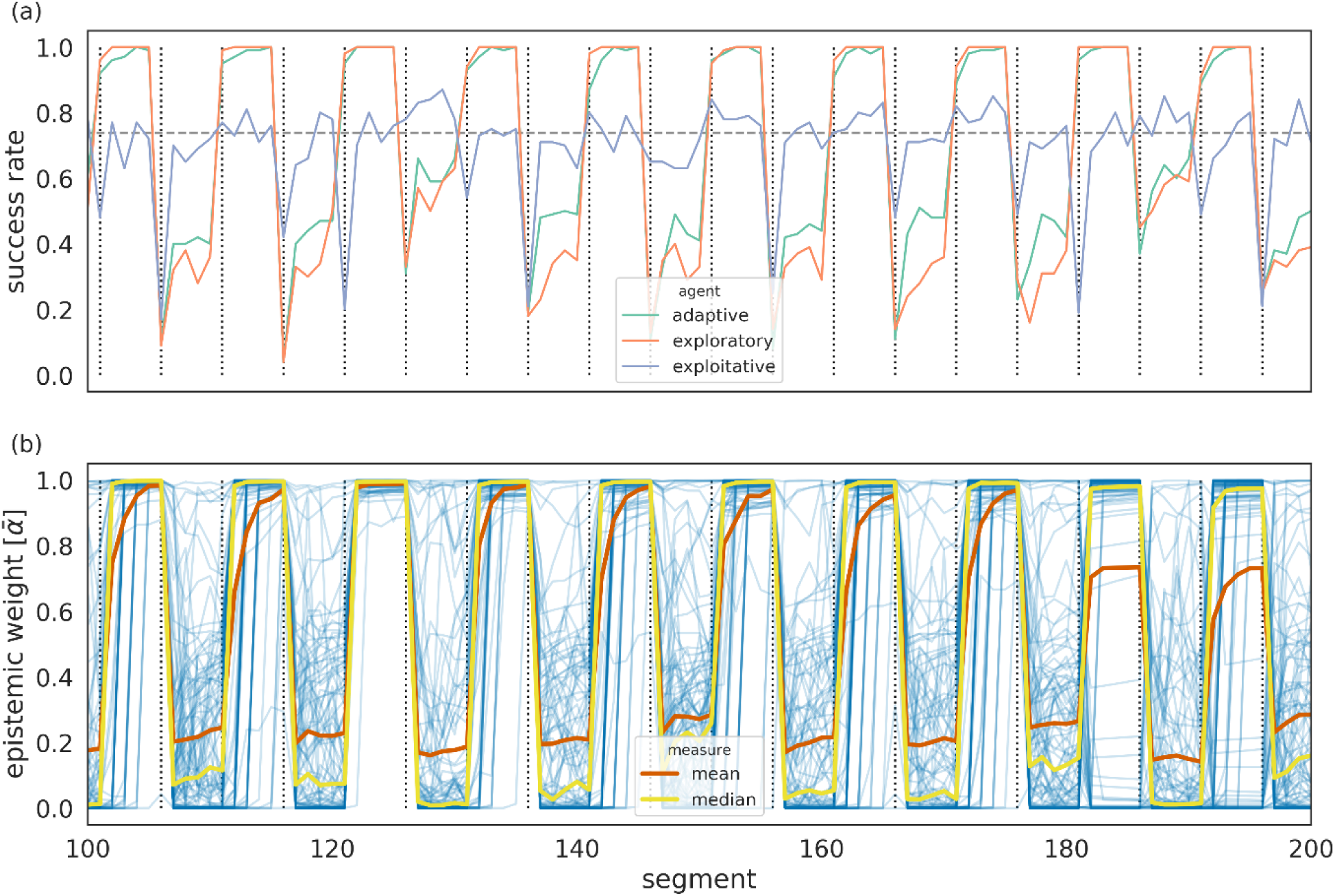
Success rates and meta-control states of an adaptive (controlled) and two non-adaptive agent types. **(a)** Group mean success rate for 100 agents of the adaptive (green), exploratory (orange) and exploitative (violet) agent type, plotted over the second half of the experiment. The horizontal black dashed line denotes the expected mean success rate for always selecting an option which returns a coloured point with probability p = 0.8. Note that the success rates of the adaptive and the exploratory agents are similar in the context variants B so that the green line is often not visible. **(b)** Trajectories of the weighting 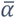 of the epistemic value contribution to the policy selection. The closer this value is to zero the more exploitative the agent becomes. To show the variability of the 100 agents’ individual 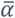 trajectories, we plotted the median 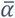 trajectory (yellow), the average 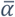 trajectory (red) and the individual 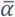 trajectories (blue). The context change schedule is the same as shown in Figure 3b.

**Figure 6.**
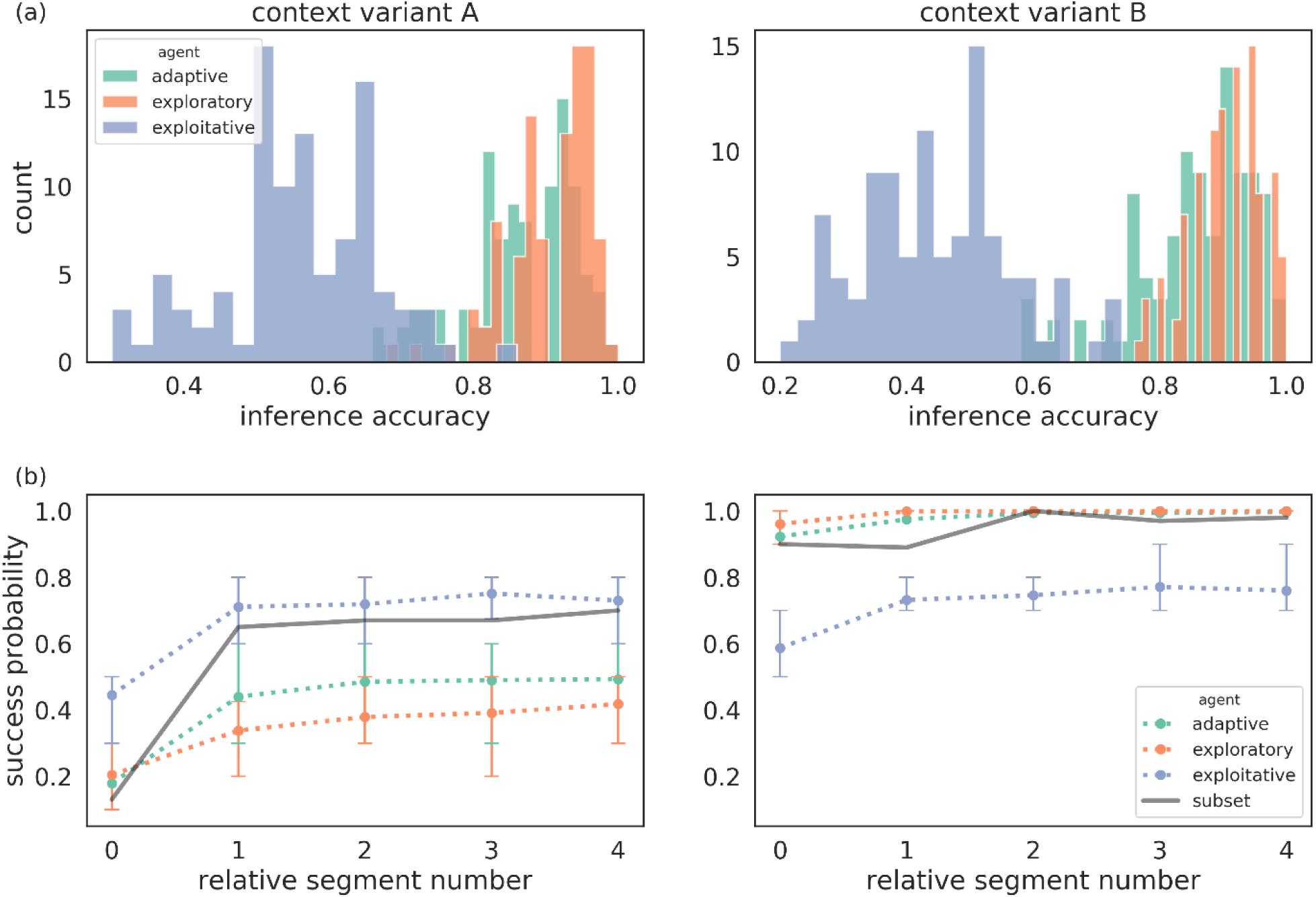
Quantification of between-agent differences in group context inference accuracy and group mean success rates. **(a)** Context inference accuracy histogram for the two contexts variants A and B, for the adaptive (green), exploratory (orange) and exploitative (violet) agent type, estimated over the last 100 segments of the experiment and defined as group probability of assigning the posterior mode to the current context. **(b)** Average success probability estimated over n = 100 instances of each agent type, over the last 100 segments of the experiment. We used the last 100 segments of the experiment to estimate success probability per instance of each agent type. The relative segment number denotes the segment number relative to the moment of context change, where zero corresponds to the segment at which the context changed. The error bars show the 25^th^ and the 75^th^ percentile. The same colour scheme as in (a) applies, where in addition, we show as black solid lines the average success probability of a subset of 10 instances of the adaptive agent which were the most efficient in down-regulating exploratory behaviour (see text for more details).

The overall low performance of the adaptive agent in the context variants A may be explained by the difficulty of downregulating exploratory tendencies in the presence of various sources of uncertainty. This is because the adaptive agent has to continuously update its beliefs about the current context, choice probabilities, and relations between the meta-control states, contexts and the success probability for a segment. In other words, the adaptive agent works as expected, but the stochasticity of its task environment keeps the adaptive agent in a limbo of uncertainty and drives the agent into an exploratory mode. This suggests that the adaptive agent can fare better in our task environment if we reduced the agent’s overall uncertainty by letting it acquire a more accurate representation of changes in their task environment. In the simulations so far, we have limited the agent to an imprecise prior on when to expect a context change, i.e. an agent expects a change after each segment with probability *p=1/5*. It is reasonable to assume that a human participant would learn after an extended period of 100 segments (500 trials) that there might be a context change around every 5 segments, where the stochasticity of the task still makes the exact duration of a context difficult to predict, but at least there should not be an anticipation that there is a context change after each segment. If we gave such a prior about the duration between context switches to an adaptive agent, it could in principle anticipate the moment of change more accurately and maintain high precision on the current context for a longer time. In the next section, we will show how representing the moment of change can improve the performance of adaptive agent and bring it much closer to the performance of the exploitative agent (IV agent) in context variants A.

### Anticipatory control of behaviour

The agents described so far were limited to expecting context change in every segment with a constant switch probability (of *p = 1/5*). Here we enable agents to represent the temporal structure of the task better and anticipate a switch around every five segments: to understand how introducing temporal representations drives anticipatory behaviour we will not consider a precise prediction of a switch after five sements, but a low uncertainty over possible durations between subsequent changes, see Methods for details. We introduce temporal expectation by extending the representational space of the adaptive agent with durations, that is, the number of segments before the next change occurs. This representation corresponds to replacing the hidden Markov framework with the hidden semi-Markov framework (see Markovic, Reiter, and Kiebel 2019).

If the adaptive agent can form predictions about the moment of change, it can use that prediction to adapt its meta-control states and any control signal a priori, before observing outcomes of the upcoming segment. To illustrate this, we show in Figure 7 prior beliefs about the meta-control state (which is represented by the weighting factor 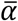) for two variants of an adaptive agent, one with weak predictions as we used in the simulations above, and one with strong predictions, see Methods for details. Importantly, one can see that the agent with strong predictions also changes its prior beliefs about its meta-control states when anticipating change (i.e., at the relative segment number 0 the group mean prior beliefs are reduced already before the change was observed in terms of outcomes). In contrast, the agent with weak predictions (i.e. the adaptive agent described above with a constant switch probability of *p = 1/5*) changes its prior beliefs only after interacting with the environment and observing a change of context at relative segment number 1.

**Figure 7.**
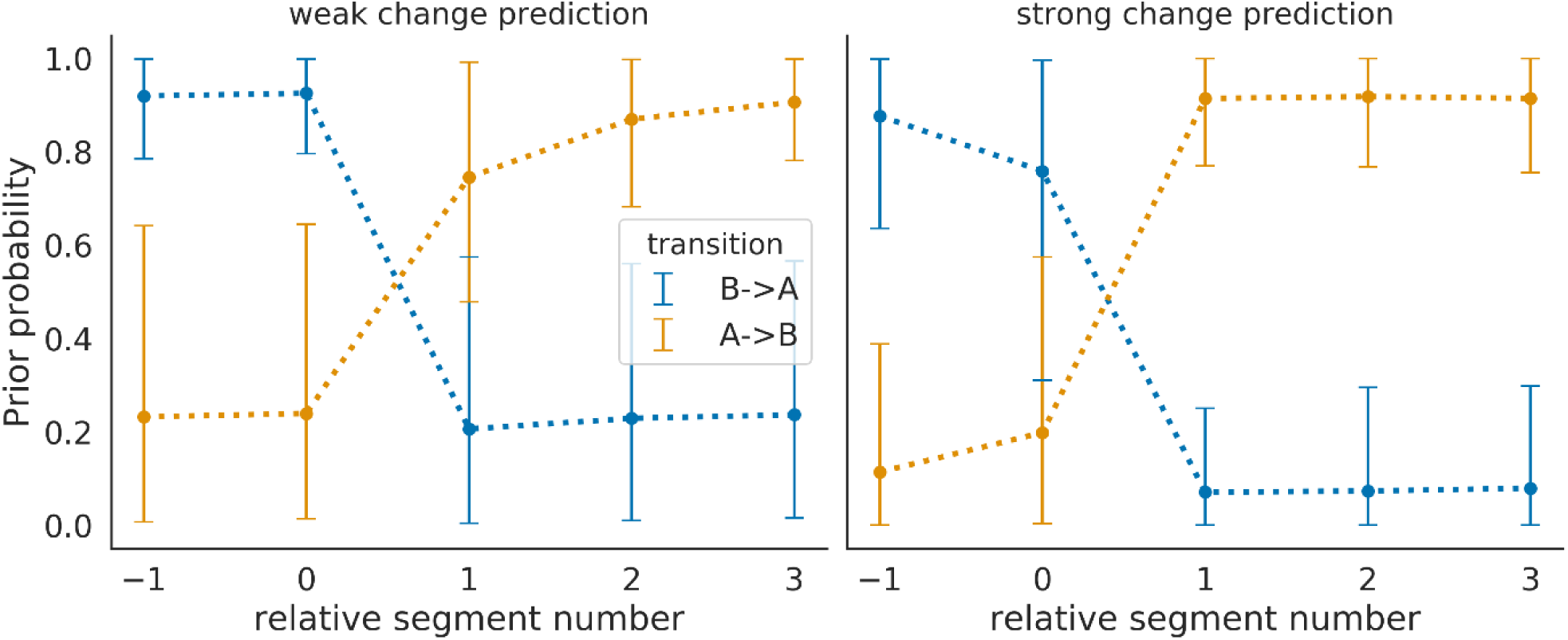
Modulation of prior beliefs over meta-control states by the anticipation of upcoming context change. **(Left)** The adaptive agent with weak change prediction, where prior probabilities over meta-control states during two types of transitions are plotted. These prior probabilities are entertained by the agent after the end of a segment before observing the outcome of the first trial of the next segment. One transition type changes from a context variant B to A (blue), the other from a context variant A to B (orange). The solid lines denote the mean, estimated over multiple transitions between two context variants, and the error bars show the 10^th^ and 90^th^ percentile. **(Right)** The adaptive agent with strong prediction, i.e. high precision on the belief about the moment of change. The agent with strong prediction, in comparison to the agent with weak prediction, adapts its prior belief over the meta-control state before having seen evidence for this change. This can be seen by comparing the prior probabilities of the two agents at relative segment number 0. One can also see that the agent with strong prediction has on average more extreme prior probabilities (closer to 0 and 1). This indicates that strong change predictions also enables the adaptive agent to gain more certainty about the current behavioural mode.

How does strong change prediction change an agent’s performance in the two context variants A and B? In Figure 8 we show a comparison of success probabilities of the three agent types. As expected, we find that all agent types benefit from strong predictions of context changes, in comparison to weak predictions, as shown in Figure 6. In context variant A, we find a significantly higher performance (p<0.05 per Wilcoxon signed-rank test) of the adaptive agent, relative to the exploratory agent, for relative segments 2 and 4. However, we expect that increasing number of instances (simulations) will trivially lead to significant differences for all comparisons. Furthermore, as the higher average performance of the adaptive agent is stable over repeated simulations (data not shown), we can exclude a chance occurance of performance differences. In contrast to adaptive agents with weak change prediction, we find that with strong change prediction the majority of agent instances (90 out of 100) down-regulates the use of epistemic value in context variants A (below 0.5 level as above). Note that the exploitative agent is insensitive to the epistemic value and therefore does not base policy selection on its subjective uncertainty about the current context. As a consequence, the exploitative agent will stick with the less informative options and have a higher chance of succeeding in context variants A. This becomes obvious for the relative segment 0 in Figure 8, where the adaptive and exploratory agents aim at reducing context uncertainty and at relative segment 4 just before another context change. Here, although the two agents have a strong prior for change prediction, they still expect the change with some probability at relative segment 4 already so that they experience increased uncertainty about their current context.

**Figure 8.**
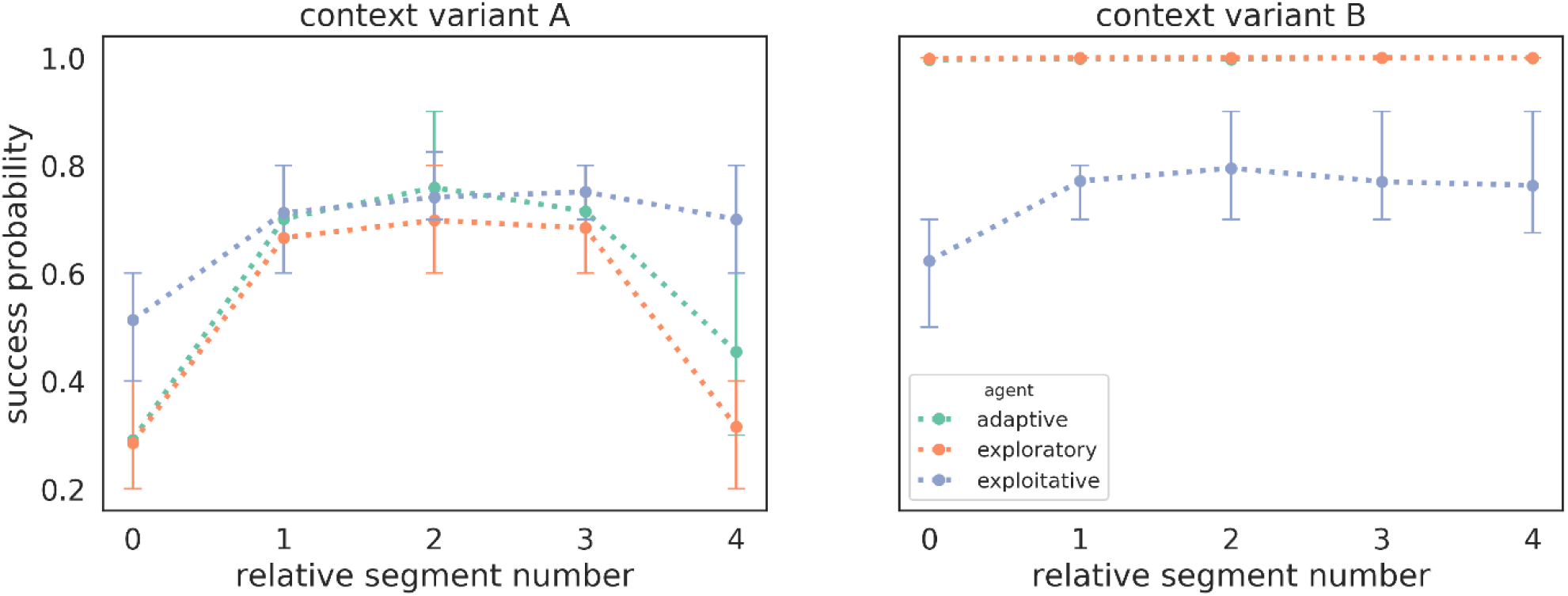
Success probability of three different agent types with strong change prediction. Mean success probability estimated as the average of success probabilities of n = 100 instances of each agent type in **(left)** context variants A and **(right)** in context variants B. Note that in context variant B the adaptive agent (green line) shows the same mean success probability as the explorative agent (orange line) so that the green line is hidden from view. We used the last 100 segments of the experiment to estimate success probability relative to the moment of change. The relative segment number denotes the segment number relative to the moment of context change, where zero corresponds to the segment at which the context changed. The error bars show the 25^th^ and the 75^th^ percentile.

Another view at the results shown in Figure 8 is to not focus on the differences in mean success probabilities, as one would in the analysis of a psychological experiment, but to evaluate agent performance from a competitive ‘survival of the fittest’ perspective. The question is then what agent type, after an initial learning period, has the highest chance to produce the best-performing agent instances, the non-adaptive or the adaptive, controlled agent? In Figure 9 we show the so-called survival function of cumulative successes of the three agent types with strong change predictions (adaptive, exploratory and exploitatory). The survival function is estimated over *n* = 100 simulations of each agent type, and as in Figure 8 we used the last 100 segments (where we pooled over context variants A and B) of the experiment to estimate success probability per instance. Critically, we found that 50% instances of the adaptive agents achieved a success probability ≥ 80%, leading to the largest probability of observing a high performing adaptive agent instance among the three agent types. For example, in an environment where an agent requires at least an 80% success probability to survive, this world would be populated mostly (66%) by adaptive agents (i.e., agents with meta-control).

**Figure 9.**
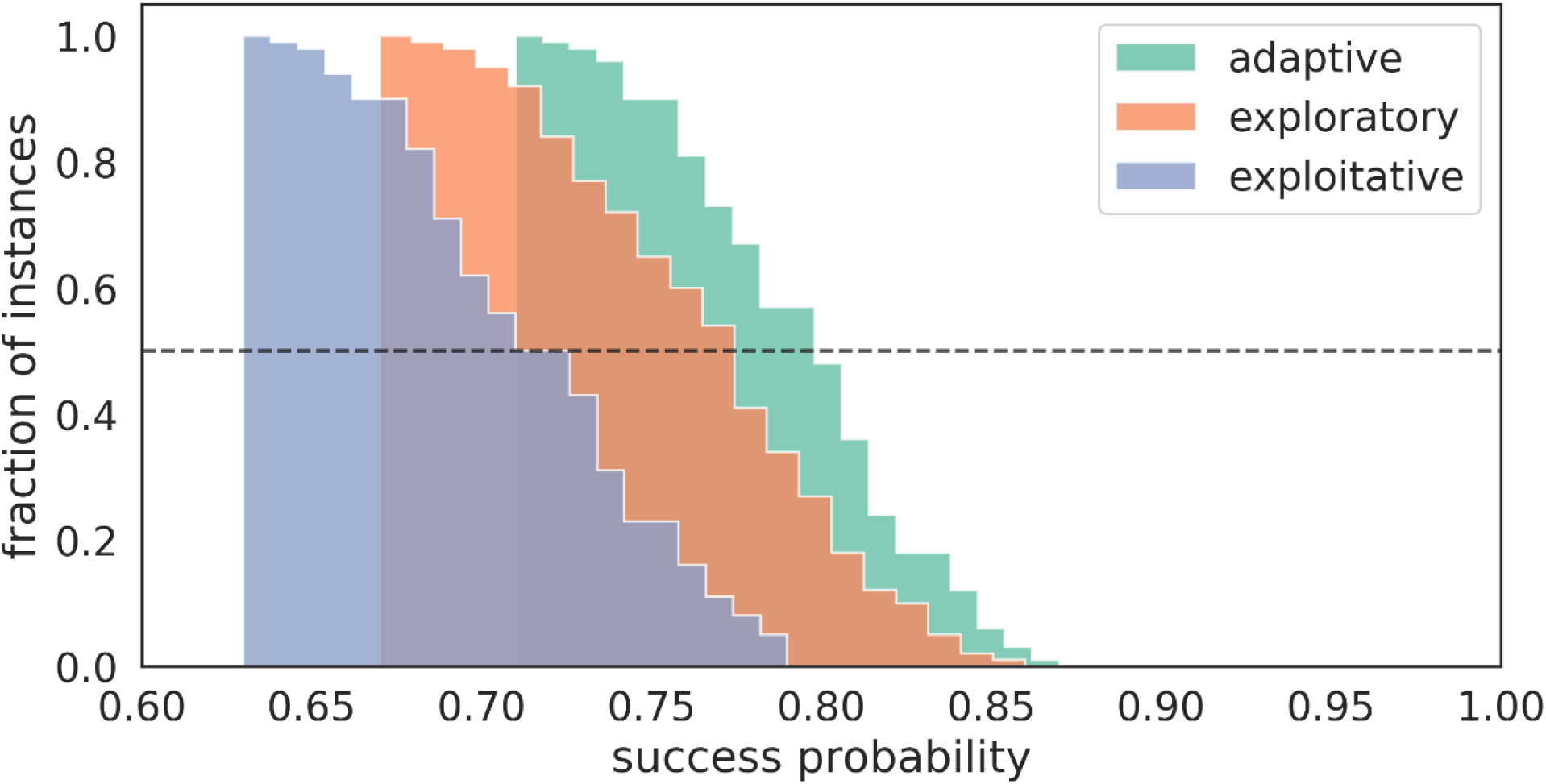
Survival function of success probabilities. Survival function (i.e., complementary cumulative distribution) for three different agent types with strong change prediction, using the same simulations as in Figure 8 over the last 100 segments of the experiment. We pooled across the two context variants A and B. The adaptive agent has the highest chance of generating a high performing instance over most success probabilities.

## Discussion

We have proposed a model which casts meta-control as an arbitrating, context-specific mechanism underlying planning and decision making under uncertainty. We used the example of the exploration-exploitation dilemma to illustrate how an agent adapts its behavioural modes (encoded as the so-called prior over policies), i.e. its internal preferences to specific sequences of actions. Critically, the agent arbitrates between explorative and exploitative behaviour by changing the relative weight of epistemic value (expected information gain) relative to the instrumental value (expected reward) when evaluating the value of different policies. As we have shown, this context-specific weighting results in adaptive transitions between explorative or exploitative behaviour, depending on the context inferred by the agent. The key element of the proposed model are meta-control states, which encode the different modes of behaviour, and can be used to learn the association between contexts and appropriate modes of behaviour. We have shown that inference over meta-control states and control signals (which make the agent behave according to its specific meta-control states) leads to adaptive meta-control as a function of the agent’s beliefs about the current context.

### Meta-control: mapping of contexts to strategies

The model describes a way to compute meta-control as a way of associating specific contexts with specific behavioural policies (modes of behaviour). Crucially, this is precisely the way Heilbronner and Hayden (2016) describe in a recent review the hypothesized function of dorsal anterior cingulate cortex (dACC). In their section ‘Mapping contexts to strategies’ they write ‘We propose, therefore, that the dACC embodies a type of storage buffer that tracks task-relevant information to guide appropriate action … .’ Clearly, this ‘storage buffer’ may translate to the beliefs over meta-control states. In addition, it is a long-standing experimental result, which Heilbronner and Hayden use to motivate their dACC hypothesis (‘mapping of context to strategies’), that dACC also represents task-relevant states. This stance is congruent with proposals that dACC is involved when switching away from the current task set (Collins and Koechlin 2012; Duverne and Koechlin 2017) or an ongoing task (Kolling et al. 2012), where the idea is that dACC does not only represent the ongoing context including task-relevant states and prior over policies but also potentially relevant alternative contexts and in particular their associated prior over policies. In the proposed model, the representation of the current and potentially relevant alternative contexts is the only way the agent can infer, when faced with uncertainty about the current context, the appropriate setting of the meta-control states. In other words, the reason why dACC seems so involved in representing task-relevant and potentially task-relevant states states may be that inference about the current context is typically not straightforward as there are several sources of uncertainty that will obscure context identity and must be routinely resolved by the brain, even in well-controlled experimental settings. It is also of note that Heilbronner and Hayden refer to ‘strategies’ and describe dACC’s function as ‘guiding action’. This is important because in the proposed model, meta-control states do not select actions directly but instead modulate the action selection process by adapting the prior over policies. This means that the prior over policies shapes viable behavioural strategies as the prior constrains the space of available policies, and supresses selection of policies that were associated with lower performance contexts.

### Control signals

Assuming that dACC guides the action selection process (Heilbronner and Hayden 2016), it is an open question what control signals are effectively sent to lower motor hierarchies like primary motor cortex? For example, Shenhav, Botvinick, and Cohen (2013) argue that the brain should compute a control signal of a specific identity (what is controlled?) and a specific intensity (how strongly?) where it is an open question how these control signals are computed and how they modulate concrete action selection in a given task. It is precisely this sort of quantitative questions that one may address using the proposed model. For example, in Figure 5b, we show the inference of the agent how much the epistemic value contributes to action selection in a specific context and specific trial. These variations directly modulate the prior over policies and can be readily interpreted as a control signal of specific identity (what policies are preferred) and intensity (how high is the prior for each policy). In other words, the proposed model and variants may be used in the future for making testable predictions how strong specific actions are preferred in a given trial, for a specific experimental sequential decision making task where participants have to plan under uncertainty, in order to reach goals.

### Relevance of meta-control for human-machine interaction

Both in psychology and cognitive neuroscience on one side and artificial intelligence on the other, there is agreement on the question what makes human behaviour so adaptive, in contrast to machines: It is the human ability to be good at meta-control, for example at deciding how to decide (Boureau, Sokol-Hessner, and Daw 2015; Gershman, Horvitz, and Tenenbaum 2015; Caccavale and Finzi 2019; Pezzulo, Rigoli, and Friston 2015). How can humans and also other animals so quickly decide and predict what strategies and what behavioural stance are a priori the most useful in a given situation? It is not unreasonable to assume that this research question will be highly relevant for any future attempts to let (heavy) machines like autonomous cars operate close to humans, in an unconstrained fashion (see e.g., Ridel et al. 2018). The reason is that human decision making, and especially rapid switches in behaviour, can presumably be best predicted, on sufficiently long time scales, if the artificial agent’s model is informed about how humans solve cognitive control dilemmas and how these solutions constrain their policies.

### Beyond exploration-exploitation: extension to other cognitive control dilemmas

The general question of meta-control, i.e. how humans infer how to make their decisions, results in a wide range of experimentally established cognitive control dilemmas. Three examples of these are (i) the goal shielding-shifting dilemma which relates to a problem a decision maker faces when pursuing a long-term goal in multi-goal settings. To reach a long-term goal, the agent has to ignore (shield) competing goals to prevent premature goal shifts (Goschke and Dreisbach 2008). However, the agent has still to be aware of the existence of alternative goals as in dynamic environment agent should be able to flexibly switch between goals and adapt behaviour to changing task demands or reward contingencies. (ii) The selection-monitoring dilemma relates to the problem a decision maker faces when deciding to pay attention to a specific part of the environment while trying to reach a goal (Goschke and Dreisbach 2008). Typically, not all available information is relevant for the task at hand, and paying attention to all of it would be detrimental for performance. However, completely ignoring currently irrelevant information would prevent the agent from noticing a crucial change in the environment and adapting its behaviour. (iii) The anticipation-discounting dilemma relates to the problem a decision maker faces when having to decide whether or not to forgo an immediate reward and wait for a delayed but potentially more substantial reward (Dai, Pleskac, and Pachur 2018; Kable 2014; Scherbaum et al. 2013). We speculate the proposed modelling approach specific to the exploration-exploitation dilemma will enable progress into determing the computations of how the brain resolves these and other meta-control dilemmas. The key conceptual idea is to build on the assumption that control dilemmas can be formulated as an inference problem over external states (contexts), internal states (meta-control states), and control signals (actions). For example, the selection-monitoring dilemma can be also understood as a hierarchical inference problem in which an agent has to decide to which aspect of the environment it should pay attention to. The probabilistic hierarchical inference would, as we have shown here, enable an agent to infer and predict that the context might change and and at the same time infer its behavioural mode which is the most appropriate for the expected context change. One of the consequences of this inference will be that the agent will use the preferred policies for this new context and, for example, infer that different states will become task-relevant, i.e. an experimenter would measure the redirection of attention to different task features.

### Metareasoning as context inference

For artificial agents, another prominent control dilemma has been subsumed under the topic of rational metareasoning, i.e. how agents can select a strategy that selects actions in time and strikes a balance between expected computational costs and expected performance (Boureau, Sokol-Hessner, and Daw 2015; Gershman, Horvitz, and Tenenbaum 2015; Lieder and Griffiths 2017). Here, an interesting research question is whether one can reduce this type of meta-control to context learning and probabilistic context inference. The idea here is that previously encountered contexts enable the agent to learn a prior over policies for this context, see (Maisto, Friston, and Pezzulo 2019) for a recent example for modelling the arbitration between habits and goal-directed control. It would be quite cumbersome for an artificial agent to predict, in an online fashion, the computational costs of the various way of how to do a task, i.e. predict and evaluate cost-benefit ratios for specific policies (Lieder and Griffiths 2017). Rather, an alternative divide-and-conquer approach with lower computational cost would be for an agent to first learn a repertoire of contexts, i.e. a discrete tiling of its environment into contextual boxes. Conjointly, as we have shown, the agent can also learn for each of these contexts a prior over policies, which can be considered the set of default behaviour of an agent in this specific context. If the brain used such a discrete contextual tiling of its environment, phenomena like maladaptive habits, where metareasoning seems short-circuited, could be at least partially explained by suboptimal context inference, as may be the case in Pavlovian to Instrumental Transfer experiments (Garbusow et al. 2014).

## Methods

### Hidden states and observables

Hidden states and observables (random variables) are depicted as circles in the factor graph shown in Figure 2. We will use *x*″ to denote hidden states at the second level of the hierarchy and *x*′ to denote hidden states at the first level. Similarly, *o*_*k*_ denotes observations (evidence) at the second level of the hierarchy, which is defined as a binary variable (success or failure), and *o*_1:*T*_ = (*o*_1_, …, *o*_*T*_) a sequence of observations at the first level of the hierarchy. At any trial *t* an observation *o*_*t*_ at the first level of the hierarchy consists of three factors:

i. point type *f*_*t*_ ∈ {0, 1}^3^,
ii. total number of points of each type *w*_*t*_ ∈ {0, …, 5}^3^,
iii. selected option *l*_*t*_ ∈ {1, …, 4}.

Hence *o*_*t*_ = (*f*_*t*_, *w*_*t*_, *l*_*t*_). Note that the point type *f*_*t*_ is expressed as a three dimensional vector (Null – (1, 0, 0), Blue – (0, 1, 0), Red - (0, 0, 1)) hence the total number of points *w*_*t*_ is obtained as

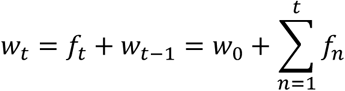

where *w*_0_ = (0, 0, 0). At the first level of the hierarchy the hidden states 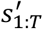consist of the following factors 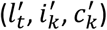, selected option, control state and context. Note that 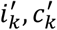 play a role of the auxiliary and constant variables at the first level, which are linked to the dynamic counterparts on the second level. The auxiliary variables are necessary to guide learning of the observation likelihood 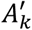, and policy selection at the first level. At the second level of the hierarchy, hidden states 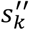 factorise into context 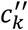, context duration 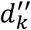, and meta-control state 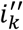, hence 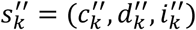.

### Likelihoods and transition probabilities

The latent state of the selected option is directly observable, hence the corresponding observation likelihood 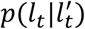 corresponds to the identity matrix. We express the relation between latent states 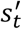 and observations *o*_*t*_ as

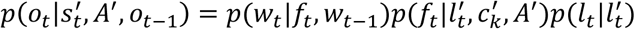

Where the likelihood over point types *f*_*t*_ is a learnable quantity

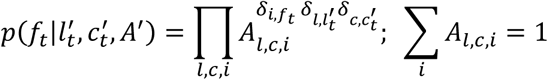

We will define the prior over point type probabilities *A*_*l,c,i*_ as a Dirichlet distribution

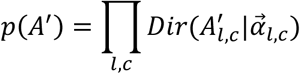

At the first level of the hierarchy policies *π*′ correspond to a sequence of five option choices, hence *π*′ = (*a*_1_, …, *a*_*T*_). Each choice deterministically sets the state of selected option 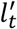, hence

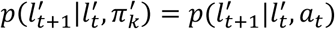

Where

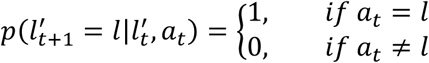

In contrast, latent factors 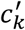, and 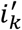 are stable during one segment. Hence their transition probabilities can be ignored.

At the second level of the hierarchy, we define the state transition probability of contexts 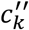 and context duration 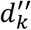 in the form of explicit duration hidden Markov model (Yu 2015), where

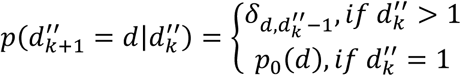

Similarly

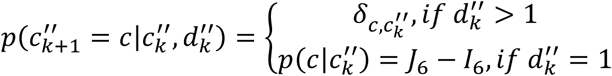

where we use *J*_6_ to denote a six dimensional all ones matrix, *I*_6_ a six dimensional identity matrix. Intuitively these state transition probabilities describe a deterministic count-down process. As long as the context duration 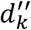 is above one, the context remains fixed 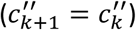 and the state duration is reduced by one 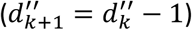. Once the duration of one is reached a new context will be uniformly selected in the next segment from the reaming five contexts, and a new context duration is sampled from the duration prior *p*_0_(*d*).

We will express here the duration prior as a discrete gamma distribution with bounded support, hence

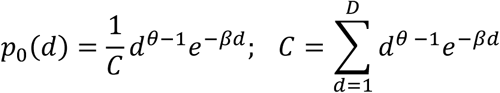

where D = 20. In **Figure 10**a we illustrate the duration priors for agents with strong (*θ* = 20, *β* = 4) and weak (*θ* = 1, *β* = 0.2) prior beliefs about the moment of change. Both priors, have the same mean but different variances. Importantly, the strong and weak priors correspond to strong and weak predictions about the future moment of change as illustrated in **Figure 10**b using an effective change probability defined as

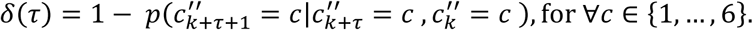

**Figure 10.**
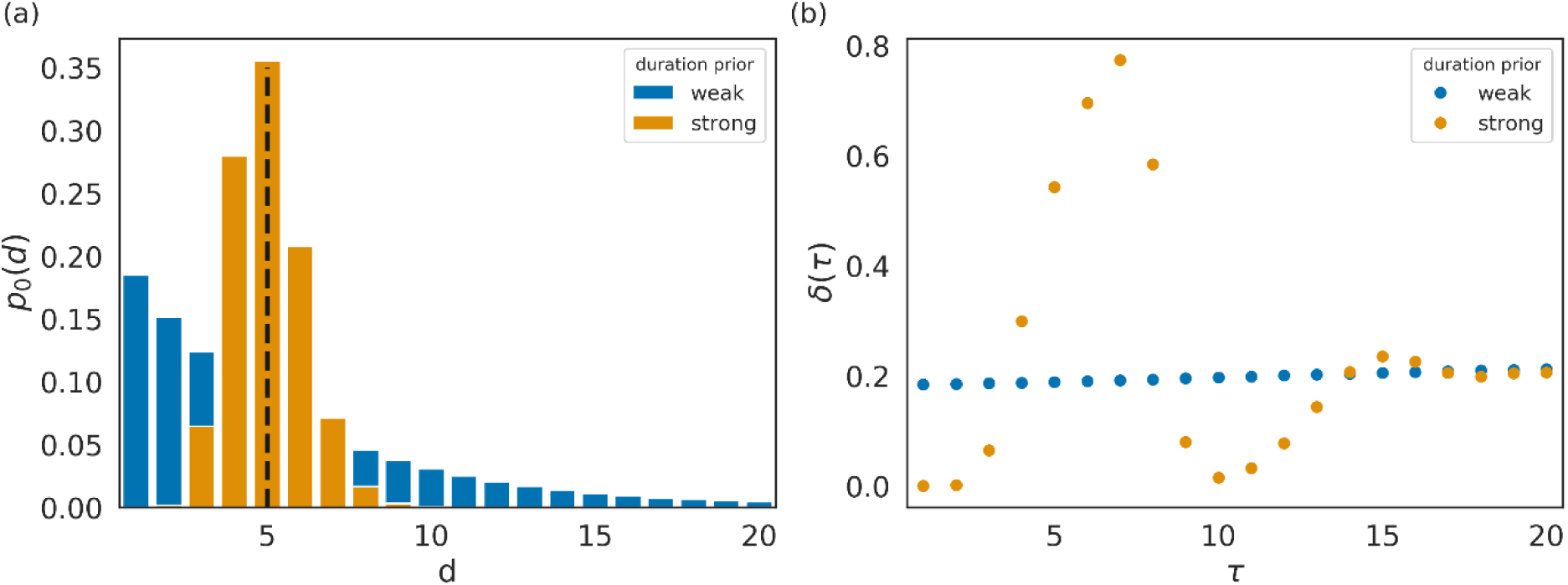
Specific cases of duration priors and the context change predictions. **(a)** Visualisation of the weak and strong prior distributions of duration d. The dashed vertical line marks the mean of both distributions. **(b)** Effective context change probability at a future segment k + τ. The effective change probability corresponds to the probability of change of current context after τ segments conditioned on a change in kth segment. Note that for strong duration prior the temporal profile of transition probability has clearly defined periods of low and high transition probability. In the case of weak duration prior the change probability δ is constant, corresponding to the hidden Markov model.

In other words, the effective change probability measures the probability that the current context *c* will change as some future segment *τ*. Note that the weak priors correspond to the hidden Markov model as the effective change probability remains constant.

### Priors over policies – expected free energy

The policy prior at different levels of the hierarchy corresponds to the expected free energy obtained as (Schwartenbeck et al. 2019)

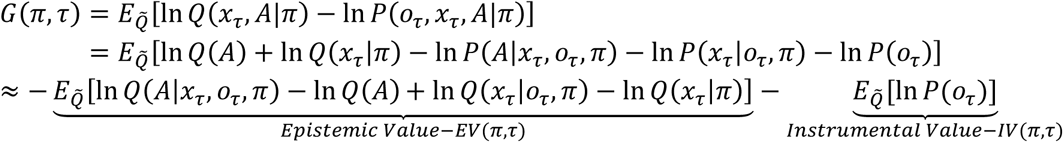

where 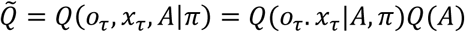 denotes a joint distribution over likelihoods *A* and prediction over states and outcomes at future step *τ* conditioned on policies *π* and likelihoods. In addition, the instrumental value (IV) term corresponds to the expectation over utility over outcomes *U*(*o*_*τ*_) = In *P*(*o*_*τ*_), where *P*(*o*_*τ*_) denotes prior outcome preferences.

In our case of a hierarchical generative model, we will adapt the above relation and define the following priors over policies and corresponding expected free energy at different levels of the hierarchy. At the second level of the hierarchy as

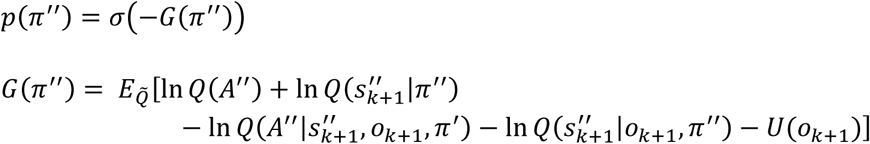

where we define the utility over outcomes as

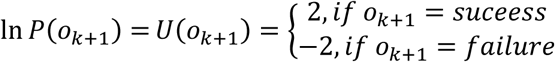

Importantly, as the expected free energy depends only on the single future step (segment) there are only two possible behavioural policies (*π*″ ∈ {1, 2}) at the second level of the hierarchy, which sets the agent either in first or second control state.

Similarly, the first level of the hierarchy as

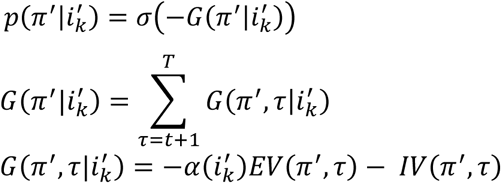

where 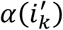 denotes a weight of the epistemic value that controls its contribution in policy selection via the second level meta-control state. Setting *α* = 1 we obtain exploratory agent variant, and setting *α* = 0 we obtain the exploitative agent variant. These two agents are non-adaptive hence they have only one available meta-control state. In contrast, the adaptive agent contains two meta-control states, hence 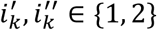 states, and the weighting function

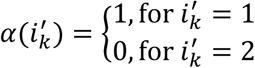

Finally, we defined the outcome utility at the first level of the hierarchy as

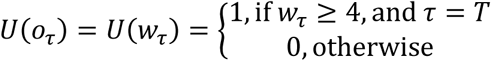

The behavioural policies at the first level of the hierarchy correspond to a set of sequences of all possible choices (option selection). Hence, *π*′ ∈ {1, …, 1024}.

### Generative model

Here we will provide a formal description of the hierarchical generative model presented in Figure 2. The joint probability distribution at different levels of the hierarchy can be expressed as

- First level

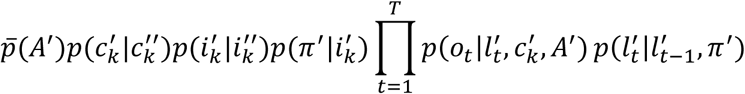

where the link distributions 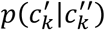 and 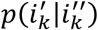 correspond to the identity matrix, meaning that for each context and control state at the second level of the hierarchy there is a matching auxiliary state at the first level of the hierarchy, and 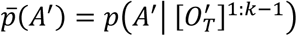 corresponds to the approximate posterior of likelihoods estimated in the last segment.
- Second level

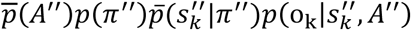

where 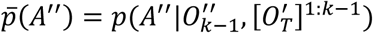 corresponds to an approximate posterior estimate at the end of the previous segment *k* − 1, and conditioned on the sequence of past observations at both levels of the hierarchy. Similarly, 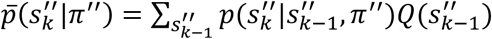, denotes the predictive probability over the current hidden states 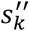.

Note that the full generative model is obtained by multiplying conditional joint distributions at different levels of the hierarchy.

### Variational inference

Inverting the generative model requires computing posterior beliefs over hidden states and behavioural policies at different levels of the hierarchy. This computation is analytically intractable and can be approximated using variational inference. Under variational inference, the true posterior is approximated as a product of multiple independent factors, hence

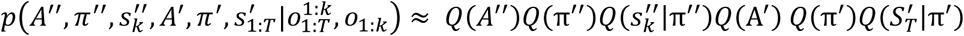

The approximate posterior is found as the minimiser of the variational free energy

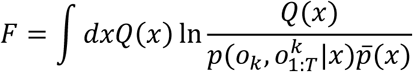

where 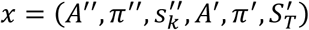. The minimum of the variational free energy corresponds to the following relations

- Second level

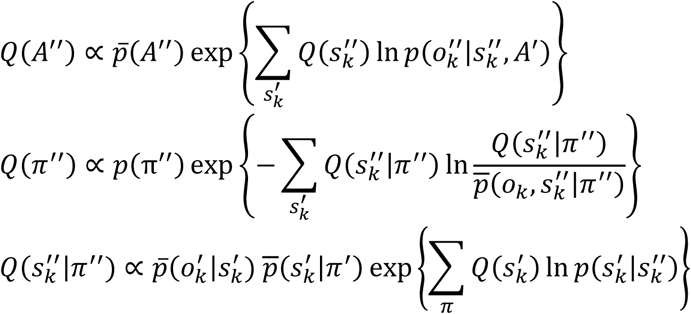

where 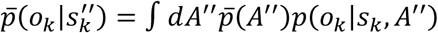.
- First level

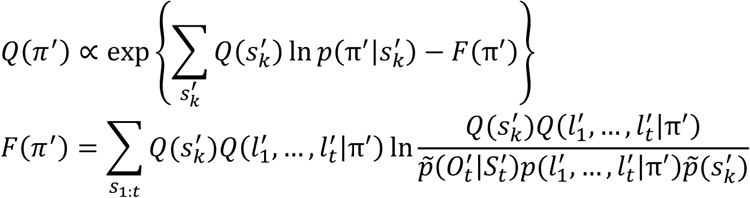

where 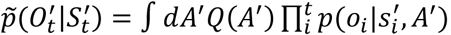, and 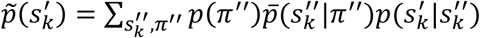.To estimate the beliefs over a sequence of hidden states 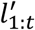 we use the Bethe approximation and the corresponding belief propagation algorithm (Schwöbel, Kiebel, and Marković 2018)

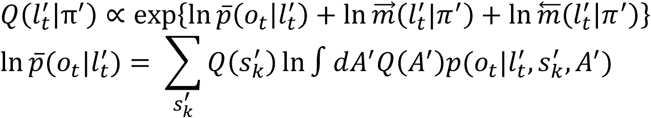 Finally, we obtain the posterior beliefs over likelihoods as

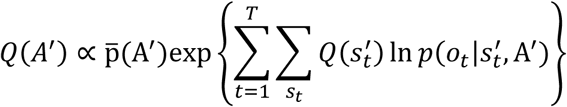

Note that we used a product of Dirichlet distributions as the prior and the posterior over likelihoods at the two levels of the hierarchy, hence we write

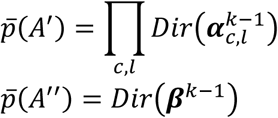

and the corresponding approximate posterior as

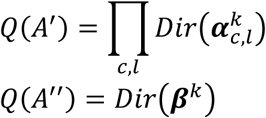

### Statistics

We use the following definitions of the group mean success rate and success probability. Lets 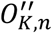 the sequence of outcomes (successes – 1, failures 0) at the second level of the hierarchy for the *n*th simulation after *K* = 200 segments. Then the group mean success rate at *k*th segment is defined as

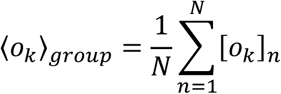

Similarly, to define instance specific success probability, we use the following relation

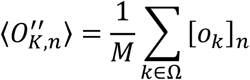

where Ω denotes set of valid segments, and *M* = |Ω|. For example, when computing success probability at different time points (relative segment numbers) of a repeated context type, the set of valid segment Ω will consist of a sequence (101, 106, …) for the relative segment number *r* = 0, of a sequence (102, 107, …), for the relative segment number *r* = 1, and so on for the three remaining relative segment numbers.

## Acknowledgments

We thank Clemens Dublaff for valuable comments and suggestions.

## Funding acknowledgments

Funded by the German Research Foundation (DFG, Deutsche Forschungsgemeinschaft), SFB 940/2, project A9 (SJK and TG), TRR 265/1, project B09 (SJK), and as part of Germany’s Excellence Strategy – EXC 2050/1 – Project ID 390696704 – Cluster of Excellence “Centre for Tactile Internet with Human-in-the-Loop” (CeTI) of Technische Universität Dresden.

## Open Practices Statement

The code and the analysis used for generating the described results is available as a github repository at https://github.com/dimarkov/pybefit/tree/master/examples/control_dilemmas.

